# Wheat *NAM* genes regulate the majority of early monocarpic senescence transcriptional changes including nitrogen remobilisation genes

**DOI:** 10.1101/2022.07.05.498473

**Authors:** Tayyaba Andleeb, Philippa Borrill

## Abstract

Senescence enables the remobilisation of nitrogen and micronutrients from vegetative tissues of wheat (*Triticum aestivum* L.) into the grain. Understanding the molecular players in this process will enable the breeding of wheat lines with tailored grain nutrient content. The *NAC* transcription factor *NAM-B1* is associated with earlier senescence and higher levels of grain protein, iron, and zinc content due to increased nutrient remobilisation. To investigate how related *NAM* genes control nitrogen remobilization at the molecular level, we carried out a comparative transcriptomic study at seven time points (3, 7, 10, 13, 19 and 26 days after anthesis) in wild type and *NAM* RNA interference (RNAi) lines with reduced *NAM* gene expression. Approximately 2.5 times more genes were differentially expressed in WT than *NAM* RNAi during this early senescence time course (6,508 vs 2,605 genes). In both genotypes, differentially expressed genes were enriched for GO terms related to photosynthesis, hormones, amino acid transport and nitrogen metabolism. However, nitrogen metabolism genes including *glutamine synthetase* (*GS1* and *GS2*), *glutamate decarboxylase* (*GAD*), *glutamate dehydrogenase* (*GDH*) and *asparagine synthetase* (*ASN1*) showed stronger or earlier differential expression in WT than in *NAM* RNAi plants, consistent with higher nitrogen remobilisation. The use of time course data identified the dynamics of *NAM*-regulated and *NAM*-independent gene expression changes during senescence, and provides an entry point to functionally characterise the pathways regulating senescence and nutrient remobilisation in wheat.

## Introduction

Wheat supplies approximately 20 percent of calories in the human diet and is an important source of protein and micronutrients. Beyond nutritional benefits, wheat grains with higher protein content are associated with increased breadmaking quality and attract a price premium. Although nitrogen (N) fertilization is commonly used to increase grain protein content, high nitrogen fertilization leads to higher production costs and environmental pollution (Aranguren et al., 2021; Martínez-Dalmau et al., 2021). Alternatively, genetic approaches can be used to increase protein content, although identifying the genetic loci to target remains a challenge.

The final grain yield and nutrient content depends on the accumulation and transport of carbon, nitrogen and other nutrients from the vegetative tissues to the developing grain. The remobilisation of nutrients is strongly influenced by the process of senescence, which is a developmentally regulated programme to remobilise nutrients from vegetative tissues to the developing grain. The starting time and progression of flag leaf senescence influences the remobilisation of nutrients and the final yield (Distelfeld et al., 2014), with the flag leaf contributing a significant proportion of nitrogen to the seed by degrading and recycling proteins (Kichey et al., 2007; Bogard et al., 2010; Havé et al., 2017). Delayed leaf senescence can be associated with prolonged photosynthesis and increased grain yield but also decrease grain protein content due to reduced nutrient remobilisation from the leaf tissues (Uauy et al., 2006; Alpuerto et al., 2021). Therefore, altering the rate and progress of senescence can influence final yield and protein content of wheat grain. Understanding the molecular components influencing flag leaf senescence and nitrogen remobilization can help to improve nitrogen remobilisation efficiency and grain protein content in wheat.

The identification of the *NAM-B1* gene which is a NAC transcription factor that influences senescence and grain nutrient content opens the door to identify the molecular pathways regulating senescence and nutrient remobilisation in wheat. *NAM-B1* was identified through positional cloning as the causal gene for *Gpc-B1* which affects grain protein content (Uauy et al., 2006). *NAM-B1*, together with its homoeologs *NAM-A1* and *NAM-D1*, influences senescence and enhance nutrient remobilisation (Avni et al., 2014; Cormier et al., 2015; Harrington et al., 2019). Most modern wheat cultivars carry a non-functional allele of *NAM-B1*, whereas the functional allele, which was identified through map-based cloning, is mainly found in wild emmer wheat and landraces (Hagenblad et al., 2012). Closely related paralogs of *NAM-B1* have been identified on chromosome 2 which also regulate senescence and nutrient remobilisation (*NAM-A2, NAM-B2* and *NAM-D2*) (Pearce et al., 2014; Borrill et al., 2019). A study of *NAM* RNAi lines with reduced expression of the *NAM-B1* homoeologs and paralogs showed that remobilisation of micronutrients and nitrogen was strongly reduced in the *NAM* RNAi lines, which directly implicates *NAM* genes in the control of nutrient remobilisation during senescence (Waters et al., 2009). These *NAM* genes provide a valuable entry point to decipher the control of monocarpic senescence and nitrogen remobilization in wheat at the molecular level.

A transcriptomic study of the same *NAM* RNAi lines at 12 days after anthesis revealed that *NAM* genes regulate transporters, hormone regulated genes and transcription factors at this early stage of senescence in flag leaves (Cantu et al., 2011). Additional *NAM*-regulated genes in flag leaves were identified by comparing wild type plants to lines mutated in either *NAM-A1* or *NAM-A1* and *NAM-B2* at 0, 12 and 22 days after anthesis (Pearce et al., 2014). Consistent with Cantu et al. (2011), *NAM*-regulated genes included photosynthesis genes and many zinc and iron transport genes. These studies provide a valuable insight into the transcriptional effects of *NAM* genes but the small number of timepoints limits our ability to understand the influence of *NAM* genes throughout monocarpic senescence. Furthermore, reduced sequencing costs and advances in genome assemblies and annotation for wheat allow more accurate analysis than was possible when previous studies on *NAM*-regulated genes were carried out using *de novo* transcriptome assemblies (Cantu et al., 2011) or earlier genome assemblies (Pearce et al., 2014; Harrington et al., 2020).

Studies using time course data can reveal the dynamics of gene expression during a developmental process. Previous studies have charaterised changes in flag leaves at the transcriptome level during senescence in wheat (Zhang et al., 2018; Borrill et al., 2019), but we do not have a full understanding of the timing of gene expression controlled by *NAM* genes for nutrient remobilization during monocarpic senescence.

To address the lack of time-resolved understanding of *NAM* gene regulation of senescence and nutrient remobilisation, we analysed flag leaf tissues at seven time points from wild type and *NAM* RNAi wheat plants. Previous work demonstrated that *NAM* genes strongly influence nitrogen remobilisation but the downstream molecular pathways were largely unknown. Therefore, we characterised gene expression changes in nitrogen-associated genes during senescence in wild type and *NAM* RNAi plants and identified genes through which *NAM* genes may influence nitrogen remobilisation. These putative *NAM* gene targets may represent target genes to improve nitrogen remobilisation in wheat.

## Methods

### Plant Material and Growth Condition

Wild type wheat (*Triticum aestivum*) plants cv. Bobwhite and sibling lines with reduced levels of *NAM* gene expression (*NAM* RNAi) were generated by Uauy et al. (2006). All plants were grown as previous described in Borrill et al. (2019) and the samples analysed in this manuscript for the wild type are a subset of those previously published in Borrill et al. (2019).

Briefly, we pre-germinated WT and *NAM* RNAi seeds on Whatman filter paper for 48 hours at 4°C, followed by 48 h at ∼20°C. These germinated seeds were then sown in trays (P40) containing a mixture of horticultural grit (15%) and fine peat (85%). We transferred individual plants to 1L square pots containing Petersfield Cereal Mix at 2 to 3 leaf stage. Plants were grown in light (16h) and dark (8h) at the temperature of 20°C and 15°C respectively. We tagged the main tiller in each pot for anthesis date, phenotyping and sample collection.

### Phenotypic Data Collection

We used SPAD-502 chlorophyll meter (Konica Minolta) to measure the flag leaf chlorophyll content at seven-time points (3, 7, 10, 13, 15, 19 and 26 days after anthesis (DAA)). At each time point, we recorded chlorophyll content from five independent plants, measuring eight locations along each flag leaf length, using only the tagged main tiller. Three out of five leaves measured for chlorophyll content were subsequently harvested for RNA extraction.

We measured grain moisture content at the same seven-time points (3, 7, 10, 13, 15, 19 and 26 DAA) at which we measured leaf chlorophyll content. From 5 independent plants, we harvested eight grains from the central spikelets (floret positions 1 and 2) from the tagged spike at each time point. We weighed fresh grains, then reweighed them after drying at 65°C for 72 hours to obtain dry weight. We calculated the percent grain moisture content from the difference in fresh and dry weight of a seed.

### Sample Collection

For RNA extraction, we harvested the flag leaf from the tagged main tiller at seven time points: 3, 7, 10, 13, 15, 19, and 26 days after anthesis (DAA) for both WT and RNAi lines. From each flag leaf we harvested the middle 3cm lengthways, to focus on a developmentally sychronised section of tissue. Three independent replicates were harvested for each timepoint and genotype. The samples were flash frozen in liquid nitrogen and stored at -80°C.

### RNA Extraction

We ground the leaf samples to a fine powder using mortar and pestles pre-chilled with liquid nitrogen. RNA was extracted using Trizol by following the manufacturer’s (ThermoFisher) protocol, with 1ml Trizol added to 100mg ground samples. Genomic DNA contamination was removed by using DNAsel (Qiagen) and samples were further cleaned through RNeasy Minikit by following instructions of the manufacturer (Qiagen).

### Library Preparation and Sequencing

Library preparation and sequencing was carried out using the same methods as described in Borrill et al., 2019. Briefly after RNA quality confirmation, the TruSeq RNA protocol v2 was used for the construction of TruSeq RNA libraries on PerkinElmer Sciclone (Illumina 15026495 Rev.F). After adaptor ligation, the libraries were size selected using Beckman Coulter XP beads (A63880). The PCR used a primer cocktail which enriched DNA fragments having adaptors at both ends. Library insert sizes was confirmed by running an aliquot of the DNA library on a PerkinElmer GX (PerkinElmer CLS760672) and concentration measured using the Tecan plate reader.

After normalization, the TruSeq RNA libraries were equimolar pooled into two final pools using Qiagen elution buffer (one pool contained WT samples, one pool contained RNAi samples). Each library pool was diluted to a 2nM concentration using sodium hydroxide (NaOH). Five μL of this solution was added to 995μL of HT1 (Illumina) to give a final concentration of 10pM. The diluted library pool (120 μL) was spiked with PhiX control v3 (1% v/v) and transferred to a 200 μL strip tube and placed on ice before loading on the Illumina cBot. The HiSeq PE Cluster Kit v3 was used to cluster the flow cell on the Illumina cBot, using the Illumina PE_Amp_Lin_Block_Hyb_V8.0 protocol. After clustering the flow cell was transferred onto the Illumina HiSeq 2000/2500 instrument. The sequencing chemistry was HiSeq SBS Kit v3 coupled with HiSeq Control Software 2.2.58 and RTA 1.18.64. Reads in bcl format were demultiplexed using the 6bp Illumina index by CASAVA 1.8, allowing for a one base-pair mismatch per library, and converted from FASTQ format by bcl2fastq.

### Transcriptome Analysis -Mapping

We pseudoaligned the samples using Kallisto v0.44.0 with default settings to the RefSeqv1.0 annotation v1.1 (Appels et al., 2018). We noticed that unexpectedly the *NAM* genes were more highly expressed in the *NAM* RNAi lines than the WT lines. Examining the read alignment we found that the transgenic RNAi construct was mapping to the *NAM* gene transcripts and artificially inflating *NAM* gene expression levels in these samples. To account for this, we substituted these regions of anomalous mapping in each of the *NAM* gene transcripts with Ns (613 to 623 bp, representing on average 29.3% of the transcript length; *TraesCS2A02G201800*.*1, TraesCS2A02G201800*.*2, TraesCS2B02G228900*.*1, TraesCS2B02G228900*.*2, TraesCS2D02G214100*.*1, TraesCS6A02G108300*.*1, TraesCS6A02G108300*.*2, TraesCS6D02G096300*.*1, TraesCS6B02G207500LC*.*1, TraesCS6B02G207500LC*.*2*). Samples were re-mapped to this masked version of the v1.1 annotation and all subsequent analysis used these re-mapped values. The masked v1.1 annotation is available at https://doi.org/10.6084/m9.figshare.20210774.v1. In total we analysed 42 samples: 3 replicates of 7 timepoints (3, 7, 10, 13, 15, 19 and 26 DAA) for 2 genotypes (WT and *NAM* RNAi). For comparison, the count and TPM (transcripts per million) of all samples were combined into one data frame by using tximport v1.0.3 (Soneson et al., 2015). All scripts used for the data analyses in this manuscript are available at https://github.com/Borrill-Lab/NAM_RNAi_Senescence and input files required to run the scripts can be found at https://doi.org/10.6084/m9.figshare.20210774.v1.

### Differential Expression Analysis

We filtered the data for further analysis to include only high confidence genes; expressed at >0.5 TPM at least in one-time point. This strategy excluded all low confidence gene models and low expressed genes from the data (Ramírez-González et al., 2018). In total 52,395 genes in WT and 52,626 genes in RNAi were expressed at >0.5 TPM. We identified genes that were differentially expressed at each timepoint by comparing WT and RNAi samples using DESeq2 v1.14.1 (Love et al., 2014). We then analysed the data using time-aware differential expression analysis software. The count expression levels of the genes expressed >0.5 TPM were rounded to the nearest integer to identify differentially expressed genes (DEGs) using ImpulseDE2 v1.10.0 (Fischer et al., 2018). For accuracy, we also identified DEGs through Gradient Tool v1.0 (Breeze et al., 2011) by using the TPM expression level of 52,395 genes in WT and 52,626 genes in RNAi on Cyverse (https://de.cyverse.org/de/) with enabled data normalization option (Merchant et al., 2016). To identify high confidence gene DEGs, we filtered to only consider genes as differentially expressed that were both identified by using ImpulseDE2 at padj <0.001 and Gradient Tool at z-score of > |2|.

### Group Patterns of Differentially Expressed Genes

We categorized the high confidence DEGs on basis of the first-time point at which they were either up or down-regulated according to Gradient Tool output for the WT and RNAi time courses separately. The Gradient Tool is based on Gaussian process regression for the identification of gene expression patterns either increasing (up-regulated) or decreasing (down-regulated) at each time point (Breeze et al., 2011). A gene that was first up-regulated at 7 DAA was placed in the “U07” group (up 7 DAA). While a gene that was first down-regulated at 7 DAA was categorized in the “D07” group (down 7 DAA). Few genes (∼2% of all DEGs) were both up-and down-regulated during either time course (3-26 DAA); these were assigned a group based on their first expression pattern with the opposite trend also indicated. For instance, a gene that showed down-regulation at 7 DAA and then up-regulated at 19 DAA was grouped as “D07U” (the second time point at which differential expression occurred was not reported in the grouping pattern). These grouping patterns for WT and RNAi are available in Supplementary Tables 1 and 2, respectively. The genes with both up- and down-regulation trends (∼2% of all DEGs) were excluded from further analyses.

### GO Term Enrichment

GO terms were only available for the RefSeqv1.0 annotation, therefore we used the same approach as Borrill et al. (2019) to transfer GO terms to the v1.1 annotation. We only transferred GO terms for genes which were >99% identical across >90% of the sequence (105,182 genes; 97.5% of all HC genes annotated in v1.1). Using GOseq v1.38.0, GO term enrichment was done for each group of DEGs separately (groups were assigned based on first-time point gene expression pattern either increasing (up-regulated) or decreasing (down-regulated)).

### Nitrogen Orthologs Identification

We identified a list of genes involved in nitrogen metabolism in Arabidopsis through a literature search (Su et al., 2004; Grallath et al., 2005; Hirner et al., 2006; Stacey et al., 2006; Masclaux-Daubresse et al., 2008; Havé et al., 2017; Gaudinier et al., 2018; Brumbarova and Ivanov, 2019). We then identified their respective orthologs in wheat using EnsemblPlants ortholog information downloaded via BioMart (Kersey et al., 2018).. Due to the evolutionary distance between Arabidopsis and wheat it was not possible to assign 1:1 orthologs in many cases due to within-lineage duplications and gene losses. Therefore, we took an inclusive approach to identifying orthologs, considering that all wheat genes in the gene tree could be orthologs of the associated Arabidopsis gene (Supplementary Table 3) Functional annotation of nitrogen associated genes differentially expressed in WT and RNAi were obtained from literature searches and g:Profiler (Raudvere et al., 2019).

## Results

### Phenotypic data and *NAM* gene expression

To examine the transcriptional differences during the initiation of senescence in wild type and plants with reduced *NAM* gene expression (NAM RNAi), we harvested an early time course of flag leaf senescence at 3, 7, 10, 13, 15, 19, and 26 DAA (Figure 1A and B). SPAD chlorophyll meter readings recorded from the flag leaves were similar from 3 to 19 DAA in both WT and RNAi, with a significantly reduced value at 26 DAA in WT compared to RNAi (Figure 1C). Grain moisture content decreased significantly between 3 and 26 DAA for both genotypes at a similar rate. By 26 DAA, the grain moisture content (55% in WT and 57% in RNAi) indicated that the wheat plants had reached soft dough stage (GS85) and the time period sampled included the majority of the grain filling period (Figure 1D; (Zadoks et al., 1974). We found that as expected, *NAM-A1* and *NAM-D1* were expressed at lower levels in the *NAM* RNA interference (RNAi) line compared to WT at same seven timepoints for which phenotypic data were recorded (Figure 1 E-F). The *NAM2* homoeologs were expressed at lower levels than *NAM1* (Figure 1E-I) with smaller differences between WT and RNAi.

**Figure 1:**
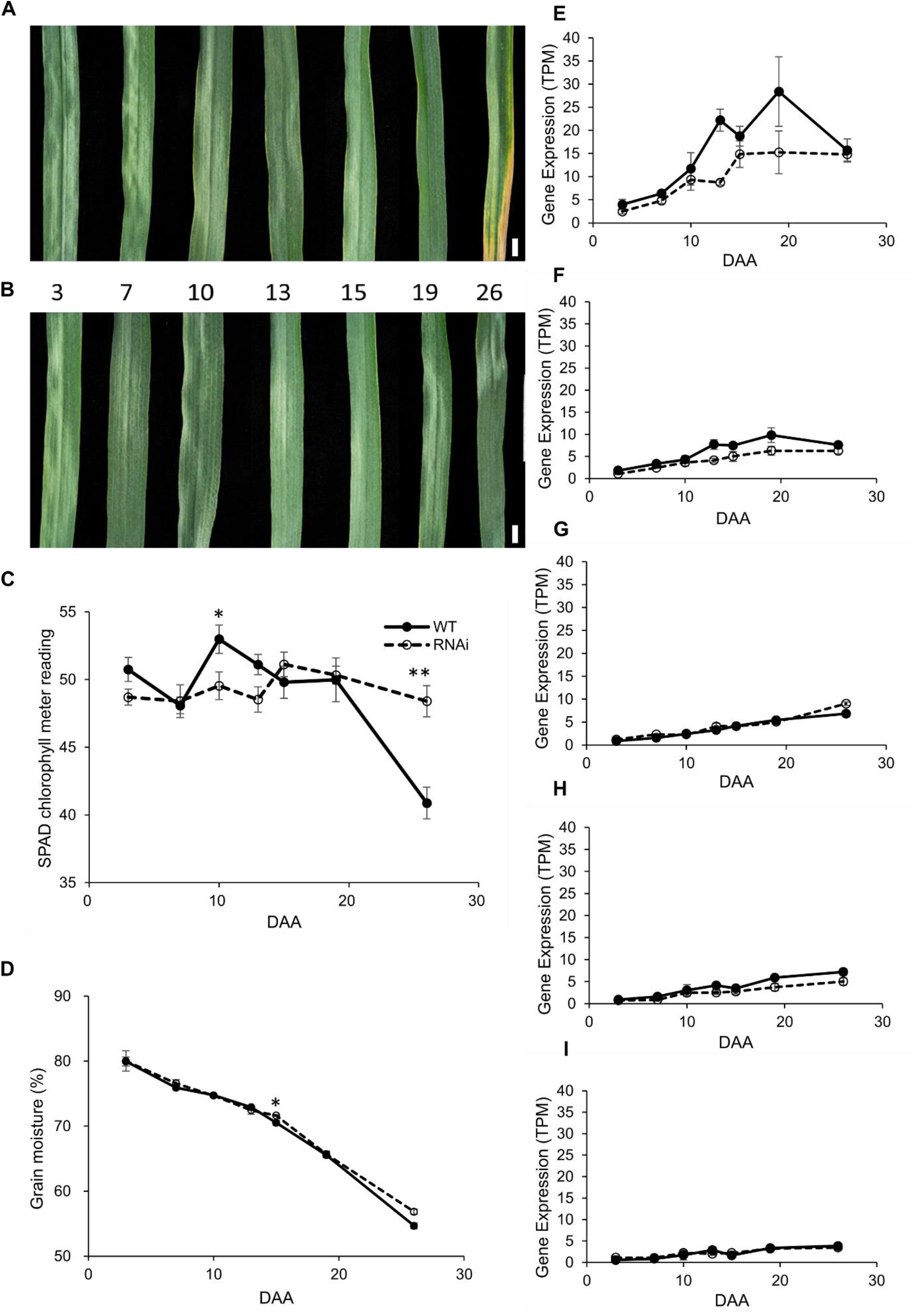
Characterization of wild type (WT) and *NAM*-RNAi plants in time course of flag leaf senescence from 3 to 26 days after anthesis (DAA). A and B) flag leaf images of WT and *NAM* RNAi from 3 to 26 DAA (WT images in A) originally published in Borrill et al., 2019), C) SPAD chlorophyll meter readings for flag leaves across the time course from 3 to 26 DAA, n=5, D) grain moisture content of grains across the time course from 3 to 26 DAA, n=5, E-I; expression pattern of *NAM-1* and *NAM-2* genes 3 to 26 DAA in WT and *NAM* RNAi measured using RNA-Seq. E) *NAM-A1 (TraesCS6A02G108300)*, F) *NAM-D1 (TraesCS6D02G096300)*, G) *NAM-A2 (TraesCS2A02G201800)*, H) *NAM-B2 (TraesCS2B02G228900)*, I) *NAM-D2* (*TraesCS2D02G214100*). Error bars represent standard error of the mean. n=5 for SPAD chlorophyll meter reading and grain moisture content and n=3 for gene expression data. Scale bar = 1 cm.

### Transcriptome Profile in WT and RNAi during senescence

#### WT plants had stronger transcriptional changes than RNAi during the time course

RNA was extracted from the flag leaf and sequenced for each of the seven time points. RNA-Seq data were aligned using kallisto (Bray et al., 2016) to the RefSeqv1.1 transcriptome annotation (Appels et al., 2018). Initially we observed artificially high levels of *NAM* gene expression in the *NAM* RNAi samples. Examining the read alignments this was caused by mapping of transcripts from the transgenic *NAM* RNAi construct to the *NAM* genes. Therefore we masked these regions of the coding sequence of the *NAM* genes with Ns to prevented artificial inflation of *NAM* gene expression in the *NAM* RNAi samples (on average 29% of the *NAM* coding sequence was masked). After re-mapping to the RefSeqv1.1 transcriptome with masked regions in the *NAM* genes, on average samples had 33.7M reads and 27.5M reads were pseudoaligned by kallisto (81.3 %) (Supplementary Table 4).

As a first step to understanding transcriptional differences between WT and RNAi we compared gene expression at each time point individually. In most timepoints < 80 genes were upregulated in WT compared to RNAi, except at 26 DAA when 549 genes were upregulated (>2-fold change, FDR <0.001; Supplementary Table 5 and 6). The 549 genes upregulated in WT at 26 DAA were enriched for GO terms associated with senescence and chlorophyll catabolism (padj<0.05; Supplementary Table 6). More genes were downregulated than upregulated at every time point, with a range from 99 to 874 downregulated genes. The largest number of downregulated genes occurred at the start and end of the time course. At the earliest timepoint 3 DAA, 693 genes were downregulated in WT compared to RNAi (>2-fold change, FDR <0.001; Supplementary Table 6) and these were enriched for GO terms associated with catabolic processes and response to freezing. At the final timepoint 874 genes were downregulated in WT compared to RNAi and these were enriched for GO terms related to photosynthesis. None of the *NAM* genes (Figure 1) were identified as differentially expressed between WT and RNAi by DESeq2, which may be due to variability between replicates and stringent p-value and fold change thresholds. Although this pairwise analysis identifies genes differentially expressed at each timepoint, it ignores information from adjacent timepoints and does not provide information on individual gene expression trajectories over the time course. Therefore we decided to identify DEGs in each genotype separately over time to reveal how dynamic gene expression is affected by the reduction in *NAM* gene expression in the RNAi lines compared to WT. We hypothesised that this approach would identify how the knock-down of *NAM* genes affects the overall senescence transcriptional programme and provide time-specific information.

To identify differently expressed genes in both WT and RNAi, we used ImpulseDE2 and Gradient Tool. We found that from 3 to 26 DAA 6,508 (WT) and 2,605 (RNAi) genes were differentially expressed. In WT out of 6,508 DEGs, 3,870 genes were upregulated and 2,638 genes were downregulated (Figure 2; Supplementary Table 1). While in RNAi, out of 2,605 DEGs, 1,585 genes were upregulated and 1,020 genes were downregulated (Figure 2; Supplementary Table *2*). During the time course, more genes were up-regulated than down-regulated in both WT and RNAi. This suggests that senescence is actively controlled through transcriptional upregulation rather than general downregulation in wheat. Approximately half of the DEGs in RNAi were also found in WT (Figure 2), contrastingly most DEGs in WT were not differentially expressed in RNAi, suggesting a unique transcriptional response in WT compared to RNAi.

**Figure 2:**
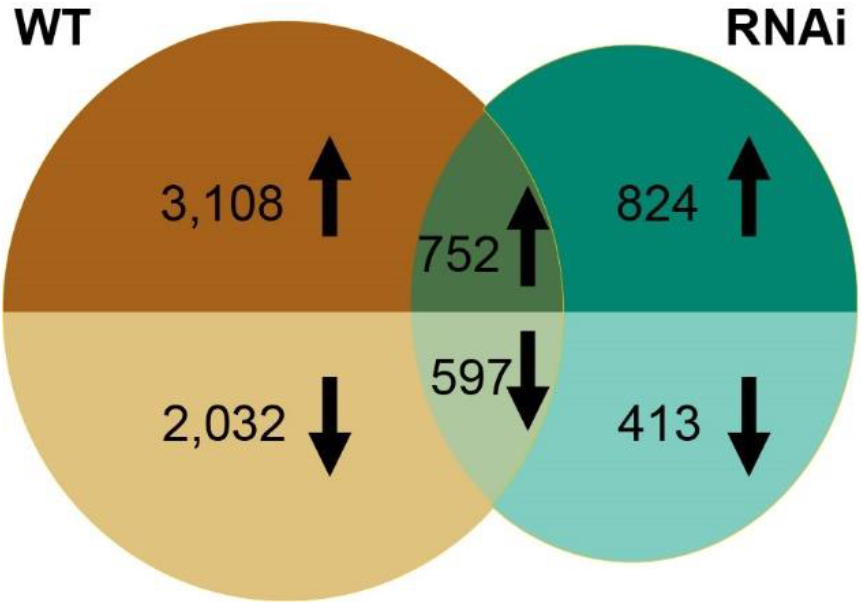
Venn diagrams of differentially expressed genes (DEGs) in WT and *NAM* RNAi. Upregulated genes are shown in the top half of each circle and downregulated genes in the bottom half of each circle. The intersection of two circles represents genes differentially expressed in both WT and RNAi. Out of 1,368 common DEGs, 19 genes were upregulated in one genotype and downregulated in the other (not shown).

#### An initial wave of downregulation is followed by upregulation of gene expression in both genotypes

To understand the temporal nature of gene expression changes, we assigned DEGs (6,508 in WT and 2,605 in RNAi) into groups according to the first time they were up-or down-regulated. For instance, a gene first up-regulated at 7 DAA would be grouped as “U07” (up 07 DAA), and a gene first showed down-regulation at this time point would be grouped as “D07”. We found that less than 2% of genes were up- and then down-regulated or vice versa during the time course in either WT (1.4%) or RNAi (1.8%) and these were excluded from further analysis. The remaining 98% of genes were described by 13 expression patterns in wild-type and RNAi (Supplementary Table 7).

In WT and RNAi, up-and down-regulation patterns were not evenly spaced over time. In both WT and RNAi, the number of genes starting to be upregulated increased during the early time points from 3 to 10 DAA, but from 13 DAA onwards the number of genes upregulated in RNAi fell to a lower level, whereas in WT 13 DAA was the timepoint with the highest number of genes upregulated (Figure 3A). Many more genes were first upregulated in WT at later timepoints than in RNAi. With the onset of chlorophyll loss at the end of the time course (26 DAA; Figure 1A), very few genes showed differential expression in either WT or *NAM* RNAi (7 genes upregulated in each line). Initiation of downregulation was stronger in the early stages of the time course in both lines, with more genes downregulated in WT than RNAi (Figure 3B). As senescence progressed, only a limited number of genes were down-regulated; 44 genes at 19 DAA in WT. In both WT and RNAi no gene was downregulated at 26 DAA suggesting that senescence process is actively regulated through transcriptional up-regulation at later stages of senescence (Figure 3A). A major shift from downregulation at the start of senescence to upregulation enduring the middle and later timepoints is evident in our dataset (Figure 3A and 3B).

**Figure 3:**
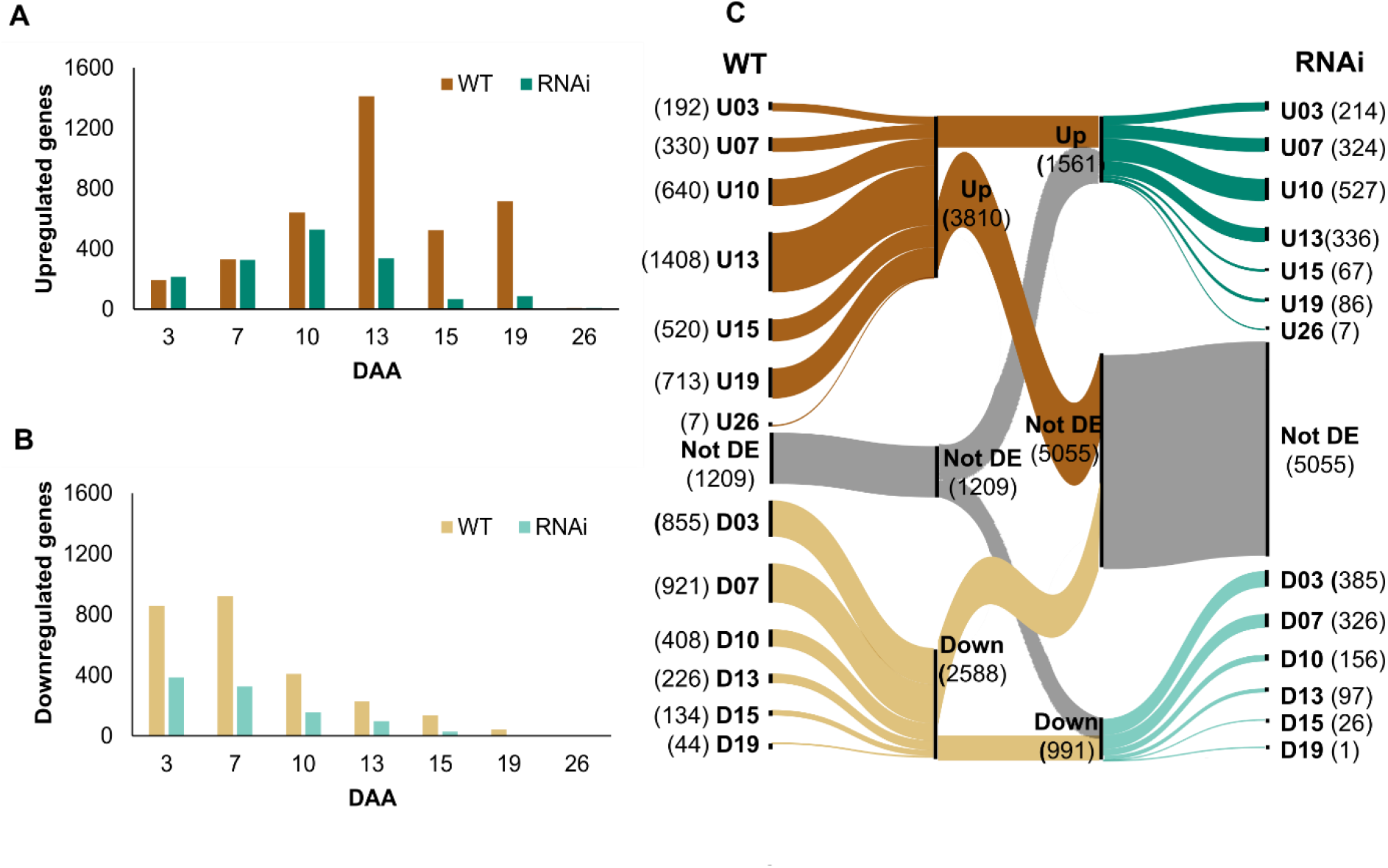
Differential expression of genes across the time course in WT and *NAM* RNAi plants. Genes are grouped according to the first time they were up-or downregulated. A) upregulated genes, B) downregulated genes, C) alluvial plot showing comparison of differential expression patterns in WT and RNAi. In C) the number in brackets for each group pattern represents number of DEGs at that time point. Not DE stands for not differentially expressed.

We found that the most of DEGs were up or down-regulated at different timepoints in WT and RNAi. For example, in WT at 3 DAA (U03) 192 genes were upregulated but in RNAi only 62 of these genes were upregulated at 3DAA while 130 of them were not differentially expressed (Not DE; Figure 3C). This limited conservation of expression profiles was common across all timepoints and in both up-and down-regulated genes (Figure 3C). We identified 1,209 genes which were not differentially expressed in WT but showed differential expression in RNAi and an even greater number were differentially expressed in WT but not differentially expressed in RNAi (5,055 genes; Figure 3C).

#### Gene Ontology (GO) term enrichments in WT and RNAi

To identify the biological processes and functions associated with each group pattern in our dataset we performed GO enrichment analysis (Figure 4; Supplementary Table 7). DEGs in WT were more strongly enriched for GO terms associated with hormones, nitrogen metabolism and other nutrient metabolism than DEGs in RNAi (Figure 4). Upregulated genes were enriched for hormone signalling and biosynthesis genes in WT but not in RNAi (Figure 4A). Up-regulated genes were enriched for GO terms associated with protein transport, proteasome, vesicle mediated transport and expressed at later time points in WT compared to RNAi (Figure 4D). Genes enriched for GO terms associated with housekeeping functions such as chloroplasts, photosynthesis, rRNA processing, and translation were downregulated at more timepoints in WT compared to RNAi (Figure 4D).

**Figure 4:**
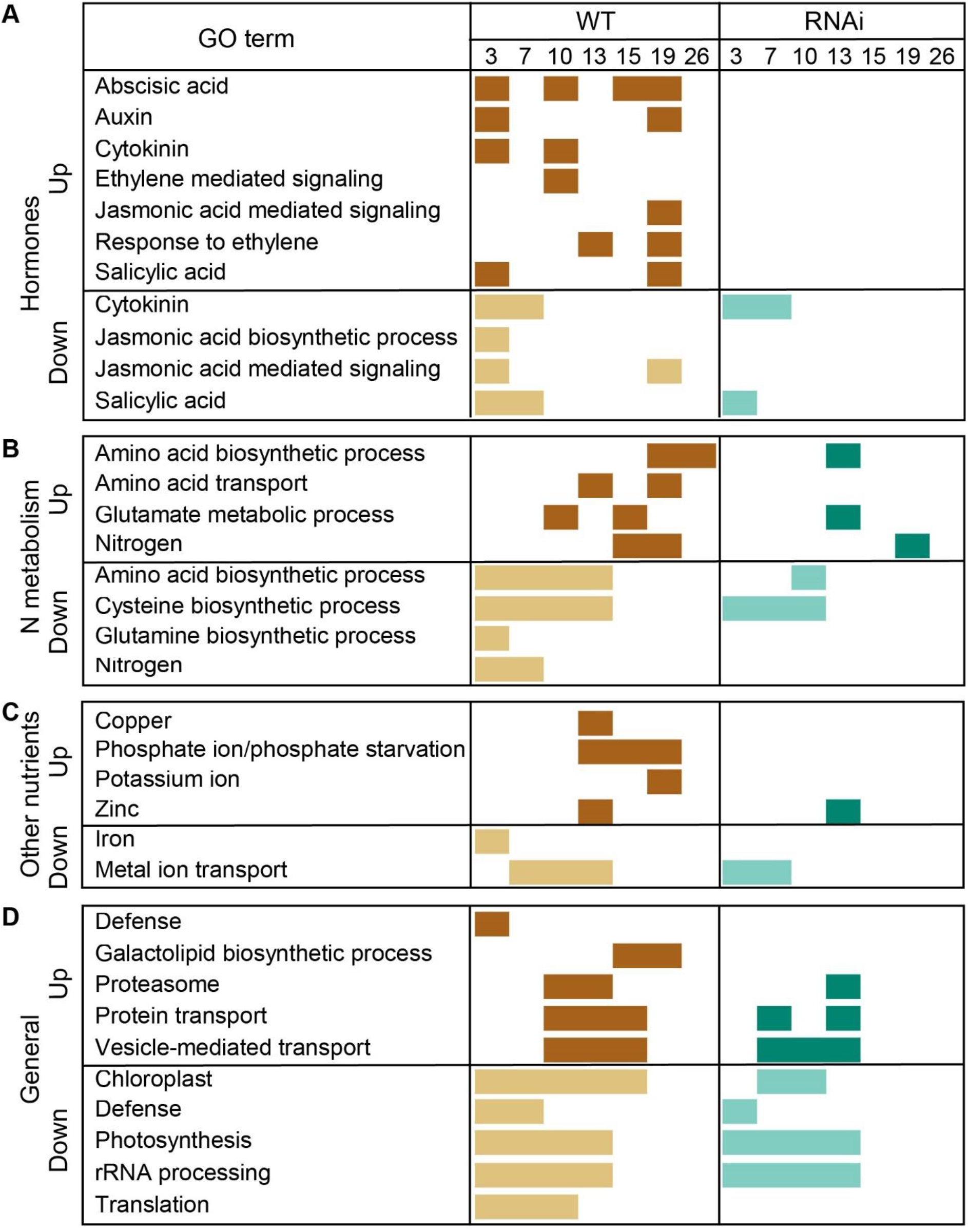
Biological processes enriched in up and down-regulated genes in wild type (WT) and RNAi lines during a time course (3-26 DAA) of senescence. Filled rectangles indicate that genes starting to be differentially expressed at that time point are enriched for that specific gene ontology (GO) term. Enriched GO terms are grouped into A) Hormones, B) Nitrogen (N) metabolism, C) Other nutrients and D) General processes. Brown rectangles represents up-regulated genes in WT; dark green represents up-regulated genes in RNAi; pale yellow rectangles represent down-regulated genes in WT and light green rectangles represent down-regulated genes in RNAi.

The differential expression patterns of genes enriched with N-associated GO terms were more obvious in WT than RNAi. GO terms related to Nitrogen (N) metabolism such as nitrogen and amino acid transport, glutamine, glutamate, cysteine biosynthesis were mostly downregulated early in the time course and then upregulated in both WT and RNAi, although the upregulation was less extensive in RNAi (Figure 4B). Genes enriched with GO terms associated with other nutrients such as copper, phosphate, potassium, and zinc showed up-regulation in WT but most of them were not enriched in RNAi except zinc at 13 DAA (Figure 4C). Genes enriched with GO terms associated with metal ion transport were down-regulated at early time points in both WT and RNAi (Figure 4C). Overall, DEGs in WT had stronger GO term enrichments, with particularly strong enrichment for processes related to hormones and nitrogen metabolism, but these enrichments were less frequently observed in RNAi.

### Genes directly involved in nitrogen metabolism

In order to identify effect of *NAM* gene on nitrogen metabolic pathway during time course of senescence, we assembled the list of genes involved in nitrogen metabolism in Arabidopsis through previous literature searches. We then identified their respective orthologs in wheat (*Triticum aestivum L*.) using *EnsemblPlants* ortholog information downloaded via BioMart. After that, we identified the expression patterns of genes involved in nitrogen transport, assimilation remobilization and transcriptional regulation in WT and RNAi lines. In total we identified 1,027 genes in wheat associated with nitrogen metabolism, of which 587 and 580 genes were expressed during flag leaf senescence in WT and RNAi, respectively. Nitrogen associated genes were differentially expressed more in WT (136) than RNAi (41) during the time course. The greater number of nitrogen associated genes DEGs in WT suggests greater changes to nitrogen remobilization or metabolism in WT than RNAi. Overall, nitrogen associated genes expressed during time course of senescence showed upregulation in WT but most of them were downregulated or not differentially expressed in *NAM* RNAi line indicating that reduced *NAM* genes affects the expression patterns of these genes in wheat.

#### Expression patterns of nitrogen transporters in WT and RNAi

During senescence, nitrogen is transported via *ammonium* (*AMT2;1*) and *nitrate* (*NRT1*.*4, NFP5*.*10, NRT2*.*5*) *transporters* across the cell membrane in the form of nitrate (NO3^-^) and ammonium (NH4^+^) ions (van der Graaff et al., 2006; Kong et al., 2016). Most nitrate transporters in our dataset were upregulated in WT flag leaves but not differentially expressed in *NAM* RNAi (Supplementary Table 3 and 8). Similarly, the highly expressed ammonium *transporter* (*AMT2;1*; *TraesCS4A02G352900*) was upregulated in WT but not differentially expressed in RNAi during our time course (Figure 5A).

**Figure 5:**
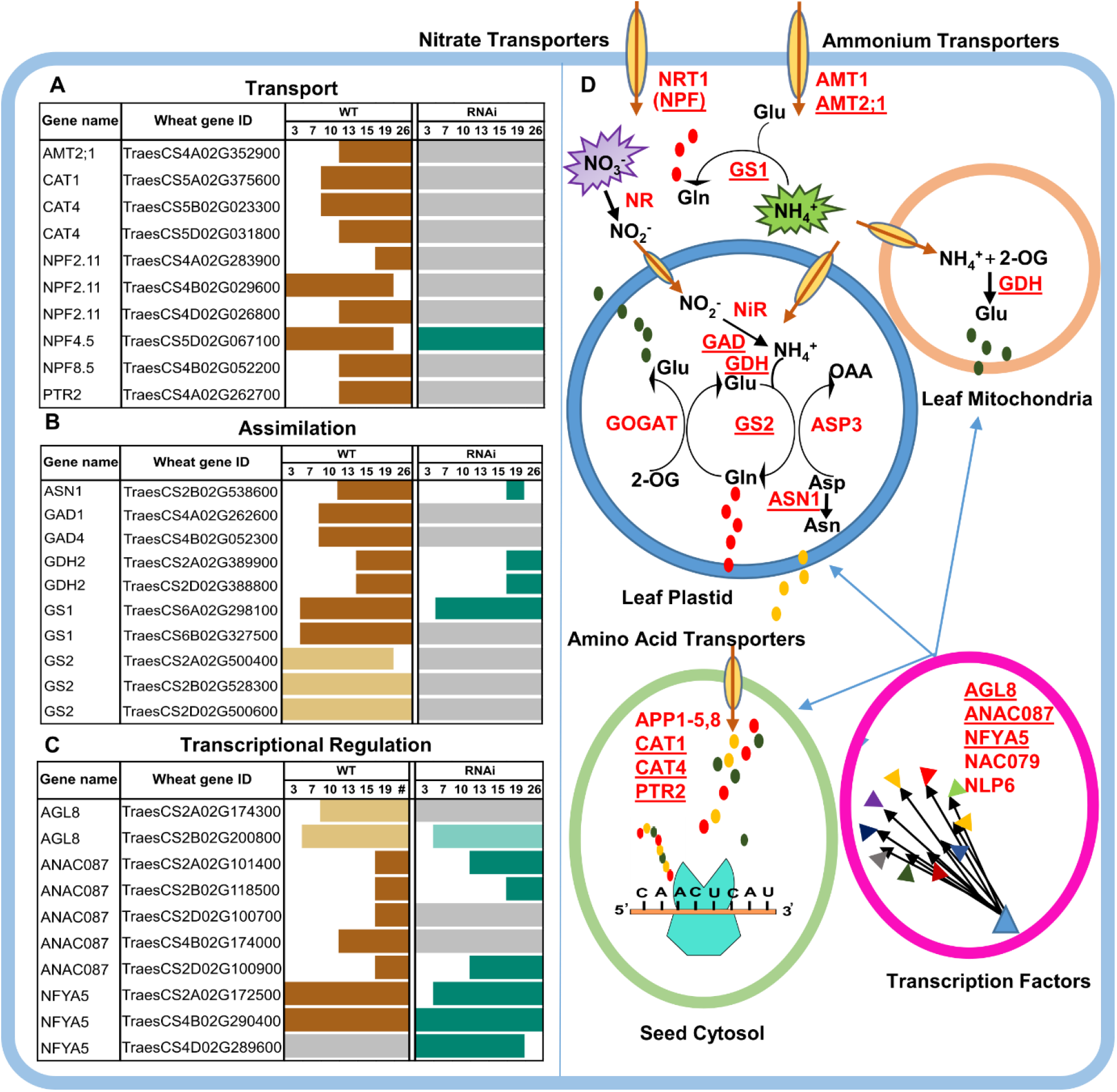
Schematic representation of genes, enzymes and processes involved in nitrogen metabolisms in wheat. The plots present on left side of figure represent differential expression pattern of the ten most highly expressed genes involved in nitrogen cycling for each category: A) transport, B) assimilation and C) transcriptional regulation. These genes are red coloured, bold and underlined in the figure to the right (D). Gene names (A-C) are given based on orthology to Arabidopsis and orthology is not always 1:1 between Arabidopsis and wheat (see Supplementary Table 3). In A-C) brown rectangles represents up-regulated genes in WT; dark green represents up-regulated genes in RNAi; pale yellow rectangles represent down-regulated genes in WT and light green rectangles represent down-regulated genes in RNAi. D) Nitrogen associated gene pathways in wheat. Ammonium (AMTs) and nitrate transporters (NRTs) transport ammonium (NH4^+^) and nitrate ions (NO3^-^) across the cell membrane. In the cytosol, Nitrate reductase (NR) enzyme reduces nitrate to nitrite. Then nitrite reductase (NiR) reduces nitrite into ammonium in the plastids. After that Glutamine synthetase (GS)/glutamine-2-oxoglutarate aminotransferase (GOGAT) cycle assimilates ammonia into N-containing compounds. Asparagine synthetase (ASN), and glutamate dehydrogenase (GDH) are involved in further assimilation of nitrogen compounds into different amino acids. Glu, glutamate; Gln, glutamine; Asn, asparagine; Asp, aspartate; 2-OG, 2-oxoglutarate; OAA, oxaloacetate. These amino acids are then transported to developing grain through different amino acid transporters (AAP, CAT1, CAT4, PTR2). All these steps are regulated by transcription factors (AGL8, ANAC087, NFYA5, NAC079, NLP6).

The deaminating activity occurring in the senescing leaf provides glutamine (Gln), glutamate (Glu) and asparagine (Asn) which are then transported to seed via amino acid transporters. These amino acid transporters include permeases (*AAPs*), proline transporters (*ProTs*), *ANT1*-like aromatic, and neutral amino acid transporters, γ-aminobutyric acid transporters (GATs) cationic amino acid transporters (*CATs*) and lysine-histidine-like transporters (*LHTs*). The amino acid transporters *CAT1* (*TraesCS5A02G375600), CAT4 (TraesCS5B02G023300, TraesCS5D02G031800), NPF2*.*11(TraesCS4A02G283900, TraesCS4B02G029600, TraesCS4D02G026800), NPF8*.*5 (TraesCS4B02G052200)* and *PTR2 (TraesCS4A02G262700)* were upregulated in WT but were not differentially expressed in RNAi (Figure 5A). Interestingly, *NPF4*.*5(TraesCS5D02G067100)* was the only amino acid transporter among ten highly expressed nitrogen transporters which was upregulated in both WT and RNAi (Figure 5A). Many other important nitrogen transporters were also expressed in our data either in WT or RNAi such as *AAP (AAP2, AAP3, AAP4* and *AAP8), PTR, CAT, GAT, LAT, LHT, ANT1* and *NPF*. Most of these amino acid transporters showed up-regulation in WT but these were either not differentially expressed or down-regulated in RNAi (Figure 5A; Supplementary Table 3 and 8).

#### Expression patterns of nitrogen assimilation genes in WT and RNAi

Many genes known to be involved in nitrogen assimilation and remobilization were expressed in our RNA seq data (Figure 5B, Supplementary Table 3 and 9) such as *nitrate reductase* (*NIA*), *nitrite reductase* (*NR*), *glutamine synthetase* (*GS*), *glutamate dehydrogenase* (*GDH*), *glutamate decarboxylase* (*GAD*) and *asparagine synthetase* (*ASN*). In general nitrogen assimilation and remobilisation related genes were more frequently up or down-regulated in the WT time course than in the RNAi time course (Figure 5B). Some genes showed later upregulation in RNAi than in WT including *ASN1 (TraesCS2B02G538600) and GDH2 (TraesCS2A02G389900, TraesCS2D02G388800)*. Other genes were up-regulated in WT but not differentially expressed in RNAi including *GAD1* (*TraesCS4A02G262600*) and *GAD4 (TraesCS4B02G052300)*. Both *GS1* homoeologs *(TraesCS6A02G298100 and TraesCS6B02G327500)* were upregulated in WT, but only the A homoeolog was upregulated in RNAi. Three homeologs of *GS2* (*TraesCS2A02G500400, TraesCS2B02G528300* and *TraesCS2D02G500600*) were down-regulated in WT but not differentially expressed in RNAi (Figure 5B).

#### Expression patterns of nitrogen transcriptional regulators in WT and RNAi

In addition to the transporters and enzymes, a number of regulatory TF genes are known in Arabidopsis to participate in nitrogen metabolism. In our dataset, the A homoeolog of *AGL8 (TraesCS2A02G174300)* was down-regulated in WT but not differentially expressed in RNAi, while its B homeolog *(TraesCS2B02G200800)* showed down-regulation in both WT and RNAi (Figure 5C and Supplementary Table 3 and 10). For *ANAC087*, the five orthologs were upregulated in WT while two of them (*TraesCS4B02G174000* and *TraesCS2D02G100700*) were not differentially expressed in RNAi (Figure 5C). Overall, we found that many more nitrogen associated genes were up or downregulated during the senescence time course in wild type than in *NAM* RNAi plants (Figure 5A-D).

## Discussion

In this study, we compared transcriptional changes in wild type and *NAM* RNAi wheat plants associated with flag leaf senescence. We found that approximately 2.5 times more genes were differentially expressed in wild type than in RNAi plants from 3 to 26 days after anthesis. Many genes associated with nitrogen metabolism are differentially expressed in wild type plants but not in RNAi plants, which is consistent with previously reported phenotypic effects of *NAM* genes on nitrogen remobilisation (Uauy et al., 2006; Waters et al., 2009).

### Dynamic transcriptional changes uncovered through time-aware differential expression analysis

The conventional approach to understand the transcriptional responses to a gene requires pairwise comparison between plants with and without the gene of interest. Using DESeq2 we carried out this pairwise analysis and identified tens to hundreds of genes differentially expressed between wild type and RNAi plants at each timepoint during senescence. Our findings were consistent with previous analyses of *NAM* RNAi and *NAM* mutant lines, including identifying changes to photosynthetic genes (Cantu et al., 2011; Pearce et al., 2014). However, specialised analysis techniques for time courses allow information to be shared between timepoints, which allows a more accurate and powerful analysis for datasets with larger numbers of timepoints. To take advantage of this we analysed transcriptional changes across our seven timepoints from 3 to 26 DAA in each genotype.

We found that although 52,395 (WT) and 52,626 (RNAi) genes were expressed in senescing flag leaves, only 6,508 (WT) and 2,605 (RNAi) genes were differentially expressed during this time period. In both genotypes, more genes were upregulated than downregulated, which shows that senescence is an actively regulated developmental process, as has been previously reported for wheat and other plant species (Breeze et al., 2011; Zhang et al., 2018; Borrill et al., 2019) Most of the genes differentially expressed in wild type plants were not differentially expressed in *NAM* RNAi plants (5,140/6,508), suggesting that *NAM* genes control approximately three-quarters of the transcription response during these early stages of senescence. We observed that WT and RNAi DEGs were split into two waves of transcriptional changes with an initial wave of downregulation followed by upregulation during later timepoints, which might not have been evident from a less time-resolved data set. *NAM* RNAi plants maintain these transcriptional waves during senescence, albeit to a lesser extent than wild type, which indicates that some transcriptional changes during senescence are *NAM*-independent, as previously proposed by Pearce et al. (2014). Nevertheless, the *NAM*-independent DEGs are much lower in number than DEGs in the wild type time course, confirming that *NAM* genes play a major role in the transcriptional regulation of early senescence in wheat (Cantu et al., 2011; Pearce et al., 2014; Harrington et al., 2020).

DEGs in WT were more strongly enriched for GO terms associated with hormones, nitrogen metabolism and other nutrient metabolism than DEGs in RNAi (Figure 4). Overall genes enriched with GO terms relating to nitrogen metabolism and nutrition showed up- and downregulation in WT but most of these genes were not differentially expressed in *NAM* RNAi. This is consistent with analysis at 12 days after anthesis which identified that genes annotated to be involved in protein metabolism and catalytic process were mostly upregulated at 12 DAA in wild type compared to *NAM* RNAi wheat (Cantu et al., 2011).

### Effect of *NAM* genes on nitrogen remobilization

Previous studies have shown that *NAM* genes affect grain protein content by altering nitrogen remobilisation in a range of genetic backgrounds and environmental conditions (Uauy et al., 2006; Waters et al., 2009; Avni et al., 2014; Pearce et al., 2014; Alhabbar et al., 2018), yet how this is mediated at the gene expression level is less well understood. To address this, we identified nitrogen metabolism associated genes in the RefSeqv1.1 gene annotation. In total we identified 1,027 genes which may be involved in nitrogen transport, assimilation remobilization or transcriptional regulation in wheat by orthology to Arabidopsis. Approximately half of these genes were expressed in our flag leaf time course in each genotype. Over three times more nitrogen associated genes were differentially expressed in WT than in RNAi across the time course (136 vs 41 genes, respectively) indicating that reduced expression of *NAM* genes affects nitrogen remobilisation at the transcriptional level during senescence. The differences in nitrogen associated gene expression between WT and RNAi may be due to direct or downstream effects of *NAM* genes which could be tested in the future using ChIP-seq or DAP-seq approaches.

We found that *NAM* genes play a significant role in controlling the expression pattern of genes associated with nitrogen transport during senescence in wheat. For example orthologs of *AAP8* (*TraesCS7B02G271151* and *TraesCS7D02G366000*) were upregulated from 10 and 13 DAA in WT, but not in RNAi. These genes had been previously shown to be highly expressed during later stages of grain development (28-30 days post anthesis; *TaAAP21*), but their potential role in the flag leaf was not noted because flag leaf samples examined were from earlier developmental stages (Wan et al., 2017; Wan et al., 2021). Manipulating these amino acid transporters has the potential to improve grain yields, nitrogen use efficiency, and protein content in crops (Dellero, 2020), and those which are *NAM* regulated (i.e. upregulated in WT but not RNAi) represent a good starting point for precise functional studies. Overall, many nitrogen transport genes were upregulated in WT but were not differentially expressed in the RNAi lines, which may indicate a true absence of transcriptional responsiveness in the RNAi line or alternatively these responses may be delayed in the RNAi line. Our analysis indicates that the widespread changes to gene expression in RNAi compared to WT are not merely a delay in timing of changes, but instead represent a loss of many transcriptional responses.

Other nitrogen associated genes showed similar trends to the transporters, with more genes differentially expressed in WT than in RNAi. For example the B homoeolog of core nitrogen assimilation gene *glutamine synthetase 1* (*GS1*) (*TraesCS6B02G327500*) was upregulated in WT but not RNAi, however the A homeolog (*TraesCS6A02G298100*) was upregulated in both WT and RNAi but to a higher maximum level in WT than RNAi. The upregulation of the A homoeolog in the RNAi as well as the WT, is consistent with *NAM* RNAi lines still being able to remobilise some nitrogen, albeit to a lower degree than WT (Waters et al., 2009) and with previous reports of the A homoeolog being more highly expressed than other homoeologs (Wei et al., 2021). We found that *glutamine synthetase 2* (*GS2*) was downregulated during senescence in WT, consistent with a previous study under high and low nitrogen (Wei et al., 2021). However, *GS2* was not differentially expressed in RNAi, which might indicate a loss of transcriptional control in the RNAi line across the nitrogen assimilation pathway, or a compensatory mechanism to increase nitrogen cycling.

We identified putative wheat orthologs of Arabidopsis transcription factors which are associated with nitrogen remobilisation. However, for this set of genes the differences between WT and RNAi at the gene expression level were weaker than for nitrogen transporters or assimilation genes, suggesting either that transcription factors controlling the nitrogen pathway are less affected by *NAM* genes, or that transcription factors regulating this process are not conserved between Arabidopsis and wheat. We previously found that NAC transcription factors which control senescence in Arabidopsis are not well conserved at the expression level in wheat during senescence (Borrill et al., 2019), therefore it seems likely that regulatory genes are also poorly conserved in nitrogen remobilisation. Combining the differentially expressed transcription factors identified in this study with transcription factors which respond to different levels of nitrogen application (Effah et al., 2022) may provide a fruitful avenue to prioritise candidate genes for functional characterisation.

## Conclusions

The use of time-aware differential expression analysis allows detailed analysis of the dynamics of gene expression during a developmental process such as monocarpic senescence. Here, we found that wild type plants undergo stronger transcriptional changes immediately after anthesis, than *NAM* RNAi lines with delayed senescence, including genes associated with nitrogen metabolism. Nevertheless, *NAM* RNAi lines do show some gene expression changes which are associated with senescence, indicating that there are *NAM*-independent pathways which regulate senescence in wheat. The list of putative *NAM*-regulated genes generated in this study provides a valuable entry point to dissect the pathways regulating senescence and nutrient translocation in wheat.

## Acknowledgements

The authors thank Cristobal Uauy for helpful discussions.

## Funding

This work was supported by the UK Biotechnology and Biological Science Research Council (BBSRC) through fellowship BB/M014045/1 to PB and the Designing Future Wheat Institute Strategic Programme (BB/P016855/1). PB acknowledges funding from the Rank Prize New Lecturer Award. TA was funded by a Commonwealth Split-Site PhD Scholarship. This research was also supported by the NBI Research Computing group through HPC resources and the University of Birmingham’s BlueBEAR HPC resources.

## Author contributions

PB conceived and designed the study with contributions from TA. PB carried out the RNA extractions and phenotyping. TA identified nitrogen-associated genes. TA and PB carried out data analysis. TA and PB wrote the manuscript and prepared figures and supplemental files.

## Data availability

RNA-seq data for RNAi samples has been deposited in the European Nucleotide Archive under project PRJEB53533. WT RNA-seq data was previously published in Borrill et al., 2019 and is available through PRJNA497810 in the European Nucleotide Archive. All other data is available within the article, supplemental files and from https://doi.org/10.6084/m9.figshare.20210774.v1. All scripts are available from https://github.com/Borrill-Lab/NAM_RNAi_Senescence.

## Notes

https://doi.org/10.6084/m9.figshare.20210774.v1

https://github.com/Borrill-Lab/NAM_RNAi_Senescence

### Competing Interest Statement

The authors have declared no competing interest.

